# Reverberation exacerbates effects of interruption on auditory spatial selective attention

**DOI:** 10.64898/2025.12.15.694434

**Authors:** Victoria Figarola, Wusheng Liang, Sahil Luthra, Christopher Brown, Barbara Shinn-Cunningham

## Abstract

Everyday listening requires focusing on one talker while ignoring competing sounds, a process challenged by reverberation and unexpected distractions. Here, we asked whether reverberation decreases effects of distractions by reducing the salience of new onsets, or compounds disruption by increasing task difficulty. Across five online experiments, participants recalled spatialized syllable streams presented with or without interrupters under pseudo-anechoic and reverberant conditions. Interrupters consistently impaired recall, especially the syllables following the interrupter. For the syllable immediately after the interruption, this effect was larger in reverberation than in anechoic conditions. These results demonstrate that distractions are especially disruptive in reverberant settings.

## 1. Introduction

Everyday listening often requires focusing on one talker while ignoring competing sounds. This requires segregating concurrent sound sources into perceptually distinct streams ^1–4^ in order to focus and maintain attention on the target stream ^5,6^. Spatial cues—especially interaural time and level differences—can facilitate stream segregation and support spatial attention to a stream (filtering out streams from other directions ^7,8^). Yet in real-world settings, reverberation often degrades the cues that listeners depend on for stream segregation and spatial perception, making the cocktail party problem even more challenging ^9^.

Reverberation is ubiquitous, from classrooms to concert venues. In reverberant environments, the ears receive a mix of direct sound and reflected energy ^10,11^. Early arriving reflections generally enhance speech perception by boosting sound levels; however, later-arriving reflections temporally smear and decorrelate what the ears receive, interfering with speech reception ^12–15^. Temporal smearing reduces the intensity transitions between loud and soft sounds, attenuates interaural level differences, introduces variability to interaural time differences, and increases both energetic and informational masking ^16–18^. Reverberation thus makes it harder for listeners to segregate speech streams, localize sources, focus attention, and encode auditory information in memory ^19^. However, listeners can partially adapt to reverberant environments with sufficient exposure, improving speech perception ^20^. Thus, reverberation imposes both perceptual challenges and opportunities for adaptation.

Sudden, salient sounds represent another challenge to attention. Novel or deviant events capture attention involuntarily, even when they are irrelevant to the task at hand ^21–23^. Such interruptions force listeners to reorient to the target—a process that requires time and which may therefore cause listeners to miss subsequent target content ^24^. Interruptions can also disrupt working memory, especially when they are unexpected ^23^. These effects highlight how vulnerable auditory attention is to distraction.

Reverberation and interruptions often co-occur in naturalistic listening environments. For example, a student in a reverberant classroom may be focusing on a lecture when a door slams. Yet little is known about how reverberation modulates the impact of such interruptions. Here, we consider two competing possibilities:

1. By smearing acoustic energy over time and increasing energetic masking across and within streams, reverberation might diminish the salience of transient sound onsets, reducing the perceptual salience of interruptions.
2. Alternatively, reverberation might compound disruption by not only interfering with selective attention but also degrading stream segregation and slowing down reorientation to the target.

To test these alternative hypotheses, we conducted five online spatial attention experiments in which listeners recalled syllables from a target stream while ignoring a distractor stream presented from a different direction. We explored performance for uninterrupted and interrupted trials in simulated environments that were either pseudo-anechoic or contained varying levels of reverberation. Across four experiments, we contrasted low and high levels of reverberation and trials that were either randomly intermingled or blocked by reverberation level. A fifth experiment with reverberated target and distractor streams contrasted a reverberant interrupter with an anechoic interrupter, which might be perceived as very close to the listener and thus more perceptually disruptive ^25^. We found a significant interaction between interruption and reverberation, with the interrupter causing a larger decrease in recall of the syllable just after the disruption in reverberant compared to anechoic conditions.

### 2. Materials & Methods

We conducted five online spatial selective attention experiments to examine how reverberation affects recall of a target syllable stream in the presence of competing speech and unexpected interruptions. All experiments followed a common design: Listeners attended to one of two spatially separated syllable streams and reported the five target syllables ^23^. The experiments differed in the degree of reverberation, trial structure, and in Experiment 5, the type of interrupter used.

### 2.1 Participants

Each experiment included 40-45 participants (Table 1), recruited via the Prolific portal or Carnegie Mellon’s online SONA platform. Studies were approved by the Institutional Review Board of Carnegie Mellon University. All participants were native English speakers with self-reported normal hearing. Participants provided informed consent and were compensated ($10/hour via Prolific or partial course credit via SONA) . Experiments were implemented online using the Gorilla platform.

**Table 1.**
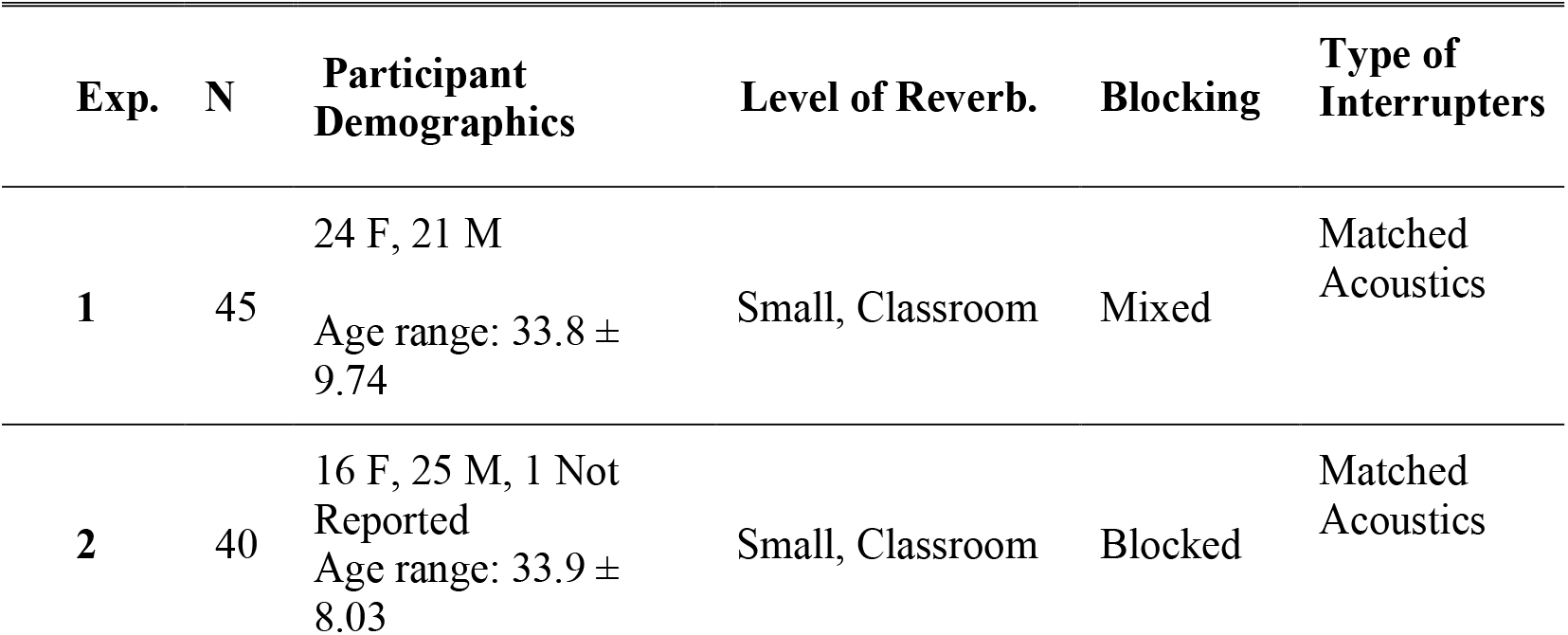
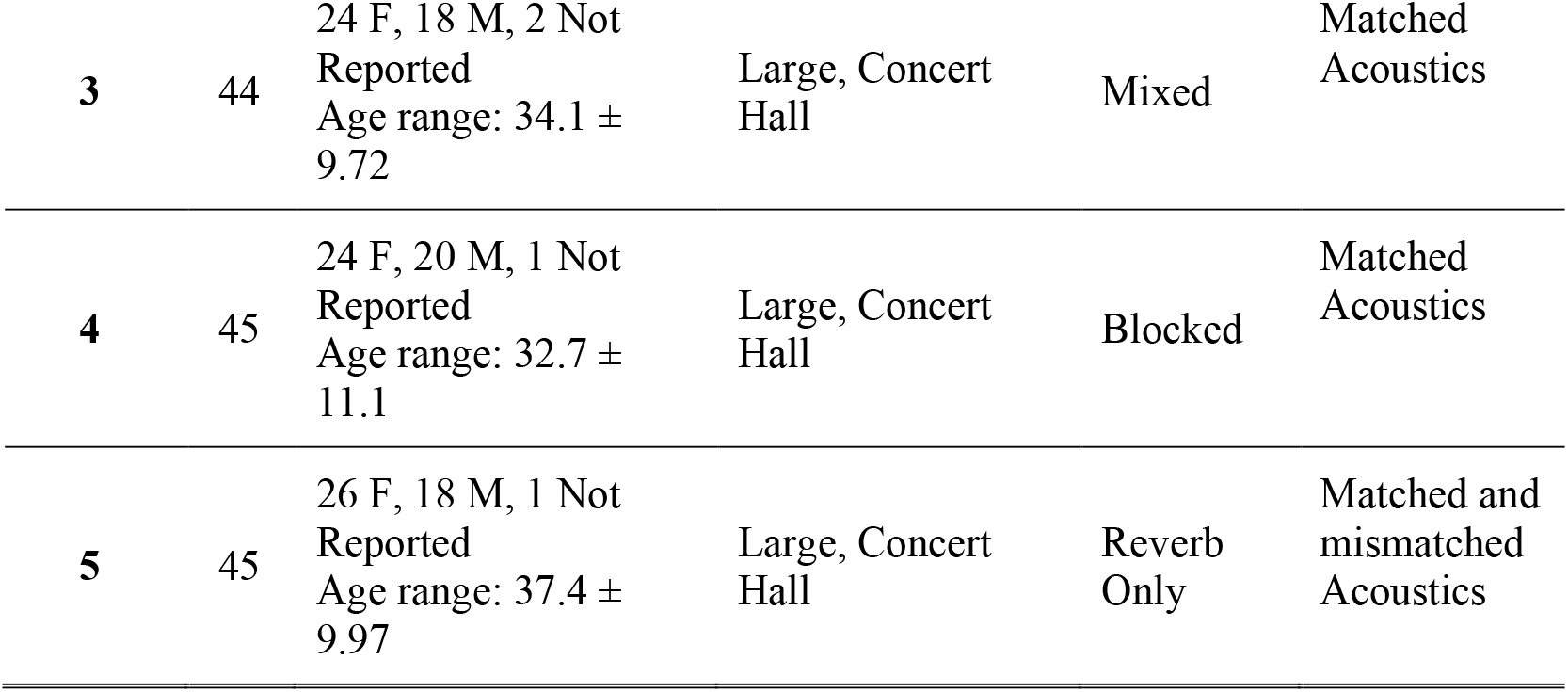
Summary of experimental design across the five studies. Each row details participant demographics, reverberation condition (classroom vs. concert hall), trial structure (mixed vs. blocked), and whether interrupters matched or mismatched the acoustic environment. Simulated reverberant environments either were a small Classroom (RT60 = 743 ms) or a large Concert Hall (RT60 = 1.91 s). N represents the number of participants.

| Exp. | N | Participant Demographics | Level of Reverb. | Blocking | Type of Interrupters |
| --- | --- | --- | --- | --- | --- |
| 1 | 45 | 24 F, 21 M<br>Age range: 33.8 ± 9.74 | Small, Classroom | Mixed | Matched Acoustics |
| 2 | 40 | 16 F, 25 M, 1 Not Reported<br>Age range: 33.9 ± 8.03 | Small, Classroom | Blocked | Matched Acoustics |
| 3 | 44 | 24 F, 18 M, 2 Not Reported<br>Age range: $34.1 \pm 9.72$ | Large, Concert Hall | Mixed | Matched Acoustics |
| 4 | 45 | 24 F, 20 M, 1 Not Reported<br>Age range: $32.7 \pm 11.1$ | Large, Concert Hall | Blocked | Matched Acoustics |
| 5 | 45 | 26 F, 18 M, 1 Not Reported<br>Age range: $37.4 \pm 9.97$ | Large, Concert Hall | Reverb Only | Matched and mismatched Acoustics |

### 2.2 Stimuli

Target and distractor streams comprised five consonant–vowel syllables (*ba, da, ga*) produced by a male native speaker of English, drawn with replacement. Streams were spatialized to 30° left or right of center. Syllables were 450 ms long with 600 ms onset-to-onset temporal spacing, and the target began 300 ms before the distractor, yielding temporally interleaved but non-overlapping sequences. On half of the trials, a novel interrupter (250 ms) occurred 125 ms before the third target syllable, spatialized to 90° contralateral to the target stream.

Across trials, environments were either pseudo-anechoic or reverberant (see 2.3 Spatialization). Experiments 1–4 included both environments, with all stimuli within a trial containing identical reverberation; Experiment 5 used reverberant streams with either anechoic or reverberant interrupters.

### 2.3 Spatialization

All syllables and interrupters were spatialized by convolving each with one of three different binaural impulse responses. Two were binaural room impulse responses (BRIRs), while the final included no reflections:

- Classroom BRIRs (Experiments 1–2): Recorded in a small classroom at Boston University (RT60 = 743 ms ^26^). A KEMAR manikin (Knowles Electronics) was positioned at the center of the room, approximately 1.5 m above the floor. A loudspeaker was placed 1 m in front of the manikin at 0° elevation. Impulse responses were recorded at azimuths from 0° to 90° in 15° increments.
- Concert Hall BRIRs (Experiments 3–5): Recorded at Carnegie Mellon University’s Fine Arts Building concert hall (RT60 = 1.91 s). The KEMAR was located at the center of the hall, approximately 1.5 m above the floor. A Yamaha MSP5A loudspeaker was placed 2 m in front of the KEMAR at 0° elevation. Microphones (Bruel & Kjaer, model 4134) were
- positioned in the ear canals. A 5-s logarithmic sweep (50 Hz–18 kHz) was presented, and responses were recorded at azimuths from 0° to 90° in 15° increments by rotating the KEMAR.
- “Pseudo-anechoic” head-related impulse responses (HRIRs; Experiments 1-5) were generated by time windowing the corresponding BRIRs to include only the direct-path sound energy and to exclude all reverberant energy ^26^.

Reverberation times (RT60) were calculated from the recorded BRIRs using the Schroeder method ^27^, implemented in the Python *psylab* package ^28^.

### 2.4 Main Task

Each participant completed 96 trials, organized into 8 blocks of 12 trials each. All experiments tested within-subject manipulations of target left vs. right and interrupted vs. uninterrupted (both factors randomized across trials within each block). Experiments 1-4 all also tested the within-subject manipulation of pseudo-anechoic vs. reverberant room conditions; in Experiments 1 and 3, this factor was randomized across trials within each block (i.e., mixed), and in Experiments 2 and 4, this factor was held constant within each block (i.e., blocked, with N/2 blocks of each type presented in random order). Experiment 5 presented only reverberant target and distractor streams and tested the within-subject manipulation of pseudo-anechoic vs. reverberant interrupter (randomized across trials within each block). Table 1 summarizes the design differences across experiments.

At trial onset, a spatialized /ba/ cue indicated the target side (30° left or right). Participants then heard the five-syllable target stream, accompanied by the interleaved distractor stream on the opposite side. At trial end, participants were prompted to recall and report the five target syllables in order.

### 2.5 Experimental Procedure

Before the main task, participants completed a headphone screening test (Huggins Pitch test ^29^) to confirm stereo listening. White noise was presented in three intervals, two of which were diotic and one containing an interaural phase difference to produce a binaural pitch percept.

Participants failing more than 4 of 6 trials were excluded. Participants then completed practice trials without interrupters and had to recall >50% of target syllables; those failing were excluded.

### 2.6 Data Analysis

Accuracy was defined as the proportion of syllables recalled in the correct position, averaged within condition per participant. We computed the effect of the interrupter by comparing target syllable recall with and without the interrupter, separately for pseudo-anechoic and reverberant conditions. Lastly, we computed the effect of reverberation by comparing target syllable recall with and without reverberation, separately for uninterrupted and interrupted conditions. All analyses were performed in R using custom scripts.

### 2.7 Statistical Analyses

All statistical analyses were conducted in R using the afex^30^, emmeans^31^, and ez packages ^32^. For Experiments 1–4, we fit repeated-measures ANOVAs with within-subject factors of syllable position (5 levels: 1–5), interruption (interrupted, uninterrupted), and environment (anechoic, reverberant). A table summarizing the results of all these analyses is provided in the supplementary materials (Table S1). Follow-up analyses used estimated marginal means (EMMs, Bonferroni corrected) to test interruption effects at each syllable position and reverberant effect by position. In addition, we conducted a separate analysis of the effect of the interrupter on syllable 3, which we expected to show the largest effect of interruption. Specifically, for Experiments 1-4, we conducted a mixed-effects ANOVA on the difference between uninterrupted and interrupted performance for syllable 3 with factors of uninterrupted vs interrupted trials (within-subject factor) and experiment.

Experiment 5 tested whether the influence of interruption in reverberant environments depended on whether the interrupter itself was reverberant (matching the room acoustics) or anechoic (as might be perceived if the interrupter were physically close to the listener). Thus, we ran a 5 (syllable position) × 3 (interrupter type: uninterrupted, anechoic interrupter, reverberant interrupter) repeated-measures ANOVA, plus a one-way ANOVA collapsing across syllables. Pairwise comparisons between conditions were conducted with Tukey adjustment.

Unless otherwise specified, significance was defined as p < 0.05 after appropriate multiple-comparison correction (pairwise EMMs, unless otherwise specified). Where appropriate, marginal effects (p < 0.10) are reported for completeness.

## 3. Results

### 3.1 Effects of Syllable Position

Across experiments, performance generally decreased with syllable position, with best performance for the first syllable (Figure 1). In Experiments 1-3, performance was slightly better for the final syllable than the penultimate syllable, while in Experiment 4, performance was roughly equal in the last two syllables. Statistical analysis supports these observations. Syllable position was significant in all experiments (all p < 0.001; Experiment 1: F(4,176) = 43.47; Experiment 2: F(4,156) = 35.99; Experiment 3: F(4,172) = 41.20; Experiment 4: F(4,176) = 44.37).

**Fig 1.**
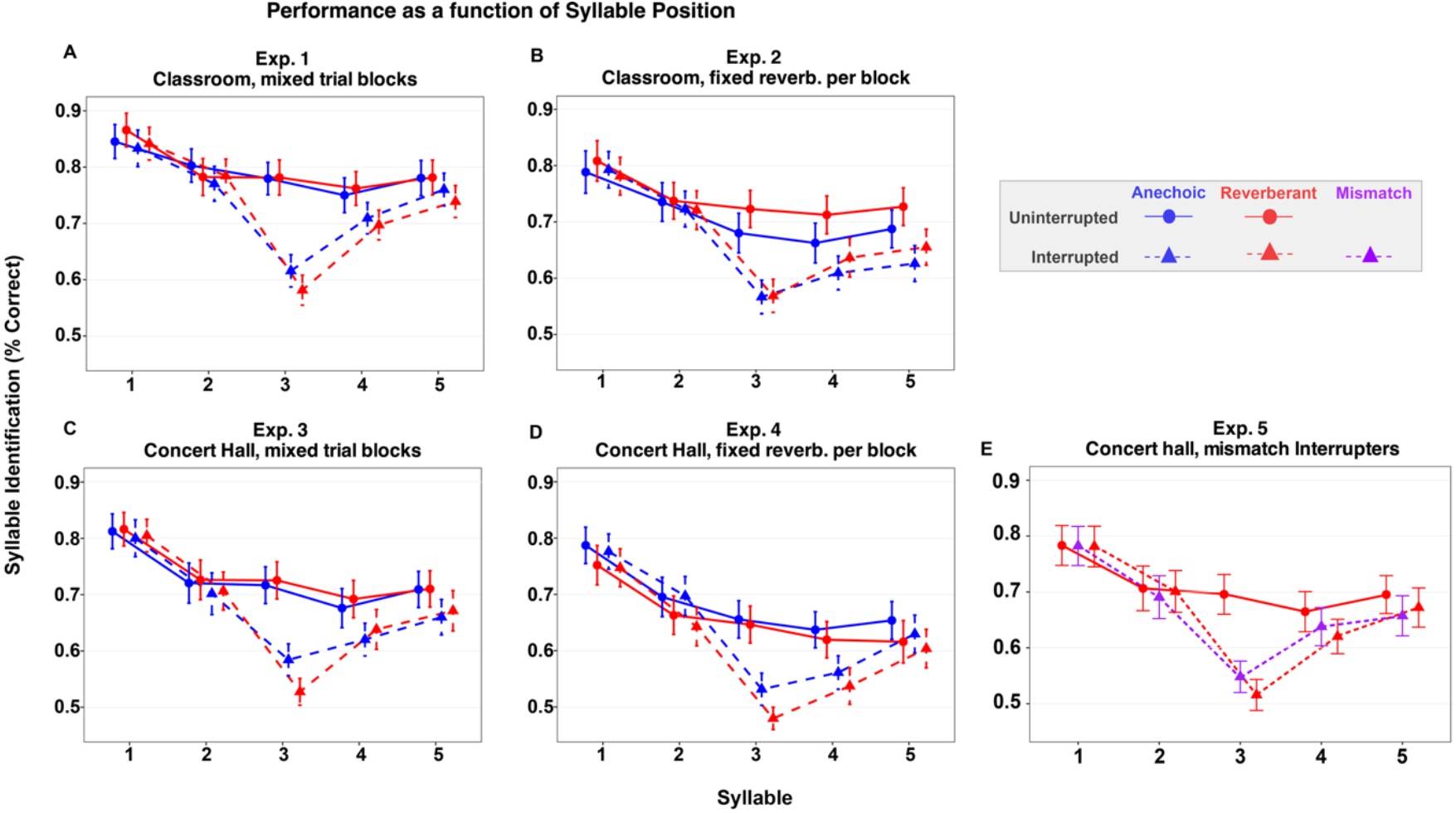
Mean syllable recall accuracy (% correct) across syllable positions for uninterrupted and interrupted trials in each experiment. Panels show (A) Experiment 1, (B) Experiment 2, (C) Experiment 3, (D) Experiment 4, and (E) Experiment 5. Circles = uninterrupted trials, triangles = interrupted trials, blue = pseudo-anechoic trials, red = reverberant trials, and purple = mismatch interrupter in Experiment 5. Error bars represent ±1 SE of the mean.

Consistent with past studies, the interrupter reduced recall accuracy differently for different syllables, with greater effects for syllables after the interrupter than those preceding it (Fig 1). Statistically, there was a significant interaction between syllable position and interruption in all four experiments (all p <0.001).

### 3.2 Effects of Reverberation

Figure 2 plots the effect of reverberation on recall, subtracting percent correct performance in reverberant trials from that in anechoic trials. In Experiment 1, there was no clear effect of reverberation (the change in performance was near zero). Experiment 3 showed a similar pattern, except for syllable 3 in interrupted trials, where performance was about 5% higher for anechoic than for reverberant trials. In Experiment 2 (classroom), performance tended to be slightly better in reverberation than in anechoic conditions, especially for later syllables (values in Fig. 2B are negative). In Experiment 4 (concert hall), performance was worse in reverberation than in anechoic conditions for all syllables (values in Fig. 2D are positive).

**Fig 2.**
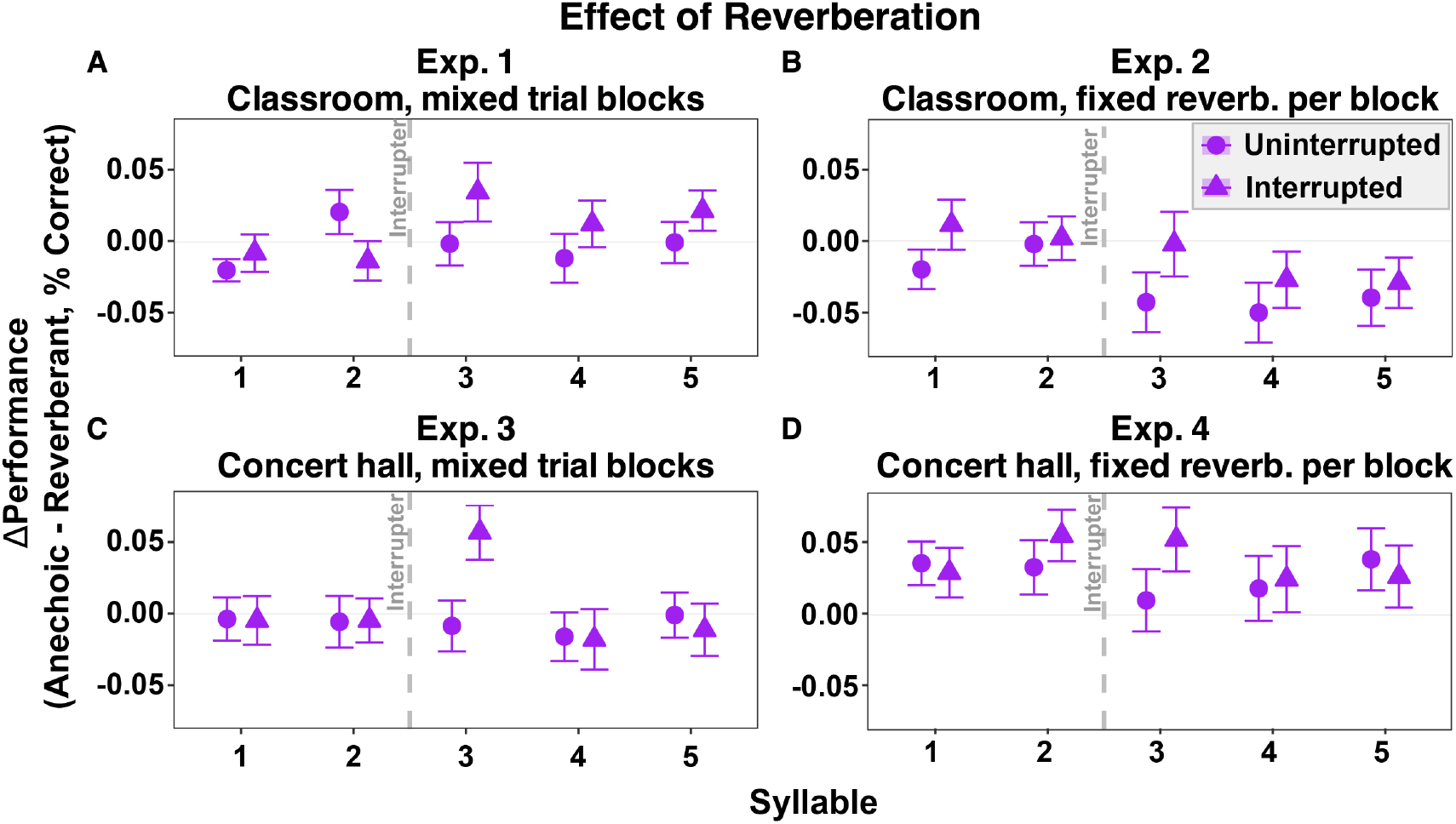
Effect of reverberation on syllable recall accuracy across experiments. Panels show (A) Experiment 1, (B) Experiment 2, (C) Experiment 3, and (D) Experiment 4. The y-axis represents the performance cost of reverberation, calculated as accuracy in the anechoic condition minus accuracy in the reverberant condition, with positive values indicating worse performance in the presence of reverberation. Circle markers indicate uninterrupted trials and triangle markers indicate interrupted trials. Error bars represent ±1 SE of the mean. The vertical dashed line marks the onset of the interrupter between syllables 2 and 3.

Statistical tests confirmed these observations. The main effect of reverberation was not significant when pseudo-anechoic and reverberant trials were intermingled (Experiments 1 and 3; p = 0.532, 0.829, respectively) but was significant when trials were blocked by room condition (Experiments 2 and 4). Specifically, accuracy was significantly higher with classroom reverberation than in anechoic conditions in Experiment 2 (F(1,39) = 8.30, p = 0.006), but significantly higher in anechoic trials than trials with concert hall reverberation in Experiment 4 (F(1,44) = 11.17, p = 0.0017).

Taken together, these results show that reverberation effects are context-dependent. In mixed designs, reverberation has little consistent impact); in contrast, in blocked designs, reverberation plays a greater role.

### 3.3 Effects of Interruption

Figure 3 plots the decrement in performance caused by the interrupter (subtracting performance in the interrupted condition from that in the uninterrupted condition). As in our previous studies, there was a clear and consistent effect of the interrupter, which was greatest for the third syllable (right after the interrupter) and persisted through syllables 4 and 5 in some cases. In all experiments, for syllable 3, the size of the interruption effect was slightly larger in reverberation than in anechoic conditions.

**Fig 3.**
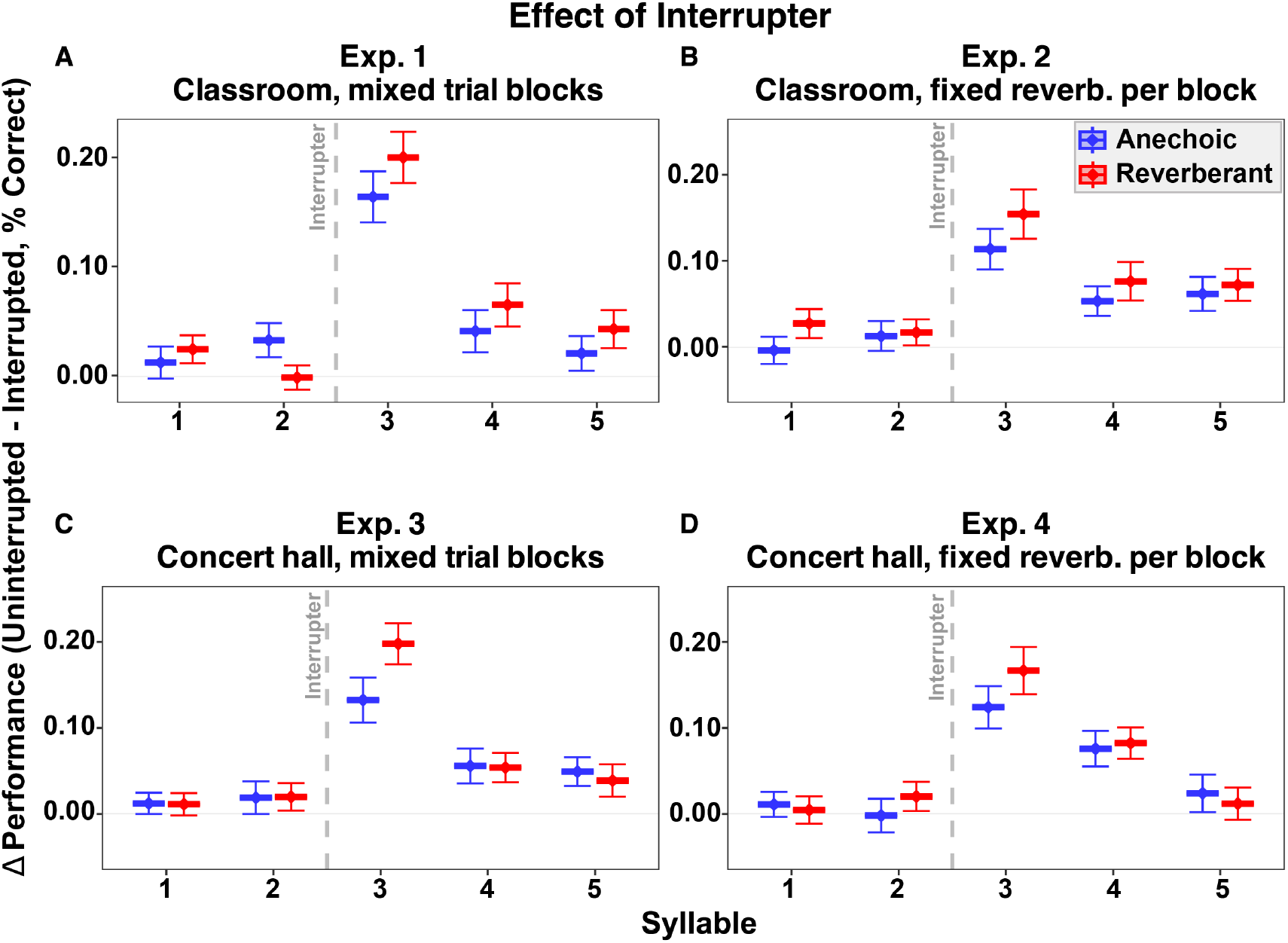
Effect of interruption on syllable recall accuracy across experiments. Panels show (A) Experiment 1, (B) Experiment 2, (C) Experiment 3, and (D) Experiment 4. The y-axis represents the performance cost of interruption (accuracy on uninterrupted trials minus accuracy on interrupted trials), with positive values indicating worse performance in the presence of an interrupter. Blue markers indicate anechoic conditions, red markers indicate reverberant conditions, and error bars represent ±1 SE of the mean.

Statistically, interrupters produced a strong main effect in all four experiments (Experiment 1: F(1,44) = 58.47, p < 0.001; Experiment 2: F(1,39) = 42.91, p < 0.001; Experiment 3: F(1,43) = 43.15, p < 0.001; Experiment 4: F(1,44) = 31.50, p < 0.001). Pairwise comparisons of syllable positions (Bonferroni corrected) confirmed that the interrupter effect was not uniform across the stream, with the largest disruptions just after the interrupter (i.e., at syllables 3 and 4; all p < 0.001). Smaller but nonetheless significant costs were also observed at syllable 5 in Experiment 1 (p = 0.022), Experiment 3 (p = 0.002), and Experiment 4 (p < 0.001), whereas syllables 1 and 2 showed no statistically significant effect in any of the experiments(all p > 0.07). Although the performance tended to be poorer than in anechoic, there was no significant main effect of reverberation or interaction between syllable position and reverberation on the interruption effect (all p>0.057).

Although the main ANOVAs showed no significant interaction between syllables, the across-experiment ANOVA examining the interrupter effect on syllable 3 revealed a significant main effect of acoustic condition (F(1,170)=10.63, p=0.001), with larger interruption costs in reverberation (mean = 0.18) than in anechoic conditions (mean = 0.13). There was no main effect of experiment (p=0.379), and no interaction between reverberation and experiment (p=0.885). Thus, reverberation reliably exacerbated the cost of the interruption on the syllable occurring just after the interrupter.

In the mismatch study (Experiment 5), the competing streams were spatialized using the concert hall BRIRs while interrupters were either anechoic (which should sound very close to the listener’s head compared to the ongoing spatialized streams) or also spatialized using the concert hall BRIRs (Fig 1E). Interrupter results for the reverberant trials were similar to those for the physically identical stimuli in Experiment 3. Performance on syllable 3 was worse with an interrupter than without an interrupter. Performance was similar for an anechoic interrupter and for a reverberant interrupter (compare blue and red values for Syllable 3 in Fig. 4). The omnibus ANOVA revealed main effects of syllable position (F(4,140) = 54.55, p < 0.001) and interrupter type (F(2,70) = 15.42, p < 0.001), as well as a syllable × interrupter type interaction (F(8,280) = 14.91, p < 0.001). Pairwise comparisons showed that both anechoic and reverberant interrupters significantly reduced accuracy relative to uninterrupted trials (Uninterrupted vs. Anechoic: p = 0.0002; Uninterrupted vs. Reverberant: p < 0.0001). However, performance did not differ between the two interrupter types (Anechoic vs. Reverberant: p = 0.881).

## 4. Discussion and Conclusion

The present study asked whether reverberation 1) weakens the salience of unexpected interrupters (reducing their disruptive impact) or 2) compounds difficulty by further challenging stream segregation. Across five online experiments, we found strong evidence for the latter.

Setting aside the effects of reverberation, the overall pattern of results confirm previous reports using a similar paradigm, but using only anechoic simulations ^23,34^: 1) Even in uninterrupted trials, performance varied with syllable position, with strong primacy effects. 2) The interrupter consistently impaired recall performance, with the largest effect on syllable 3 (which occurred immediately after the interrupter), and a smaller but persistent effect that lasted through syllables 4 and 5.

In addition to the overall effect of reverberation level, the effects of reverberation also depended on whether reverberant trials were blocked or intermingled with anechoic trials.

Specifically, performance on anechoic and reverberant trials was very similar when these trials were intermingled (Experiments 1 and 3) for both levels of reverberation (simulated classroom and concert hall), with one exception in Experiment 3 (concert hall), where the effect of reverberation on recall of syllable 3 was large for interrupted trials but not uninterrupted trials (Figure 2, panel C), hinting at a significant interaction of reverberation and interruption. When trials were blocked and reverberation was modest (classroom), performance was consistently slightly better in reverberant blocks than anechoic blocks. This suggests that the modest reverberation improved speech perception when listeners had the opportunity to adapt to the reverberation, consistent with the beneficial effects of early reflections on speech intelligibility. In contrast, when trials were blocked and reverberation was pronounced (concert hall), performance was better in anechoic than reverberant blocks.

Critically, looking across the first four experiments, the interrupter had a consistently larger effect on recall of syllable 3 in reverberant conditions than in anechoic conditions. Our results are inconsistent with the idea that the “smearing” of the acoustic scene makes unexpected events less salient; instead, in a reverberant environment, where acoustic cues for grouping are degraded and auditory scene analysis is less robust, salient events are especially disruptive to attention.

Finally, Experiment 5 showed that when listeners were focusing attention on streams with heavy reverberation, both anechoic and matching-reverberant interrupters had similar effects. We hypothesized that during a reverberant trial, the anechoic interrupter might seem particularly close to the listener, and thus especially distracting; instead, we found that the salience of anechoic and reverberant interrupters was similar.

Future work could incorporate physiological measures such as pupillometry to help test whether reverberation imposes additional listening effort and how the influence on interruption on listening effort may be influenced by room acoustics.

## Acknowledgments

Research reported in this publication was supported by the National Institute On Deafness And Other Communication Disorders of the National Institutes of Health (T32DC011499, R21-DC018408, R01-DC019126, and R01-DC022699) and the Office of Naval Research (N00014-19-12332, N00014-20-12709, and N00014-23-12065).

## Author Declarations

Authors declare no conflicts of interests.

## Ethics Approval

This study was reviewed and approved by Carnegie Mellon University’s Institutional Review Board. The participants provided their written informed consent to participate prior to their participation in this study.

## Data Availability

The raw data supporting the conclusions of this article will be made available by the authors, without undue reservation.

