## Supplemental Table 1 for "Reverberation exacerbates effects of interruption on auditory spatial selective attention"

**Supplementary Material**

**Table S1. Summary of omnibus repeated-measures ANOVAs for Experiments 1–4.** Each experiment tested a 2 × 2 design with factors of syllable position (5 levels), interruption (interrupted vs. uninterrupted), and environment (anechoic vs. reverberant). The table reports whether each main effect or interaction was significant, with p-values shown where applicable. Simulated reverberant environments either were a small Classroom (RT60 = 743 ms) or a large Concert Hall (RT60 = 1.91 s).

| | | | Statistically Significant Effect? | | | | | |
|---|---|---|---|---|---|---|---|---|
| **Exp.** | **Reverb.** | **Blocking** | **Syllable Position?** | **Int.** | **Syllable Position x Int.** | **Reverb.** | **Reverb x Int.?** | **Reverb x Syllable** |
| **1** | Classroom | Mixed | Yes (p<0.001) | Yes (p<0.001) | Yes (p<0.001) | No (p-0.53) | No (p=0.315) | No (p=0.217) |
| **2** | Classroom | Blocked | Yes (p<0.001) | Yes (p<0.001) | Yes (p<0.001) | Yes (p=0.006) | No (p=0.119) | No (p=0.147) |
| **3** | Concert Hall | Mixed | Yes (p<0.001) | Yes (p<0.001) | Yes (p<0.001) | No (p=0.82) | No (p=0.515) | No (p=0.108) |

| | | | | | | | | |
|---|---|---|---|---|---|---|---|---|
| **4** | Concert Hall | Blocked | Yes (p<0.001) | Yes (p<0.001) | Yes (p<0.001) | Yes (p=0.0017) | No (p=0.425) | No (p=0.833) |
| **5** | Concert Hall | Blocked | Yes (p<0.001) | Yes (p<0.001) | Yes (p<0.001) | -- | -- | -- |
